## Supplementary figures and images for "Cryo-EM structures reveal two distinct conformational states in a picornavirus cell entry intermediate"

### Figure S1

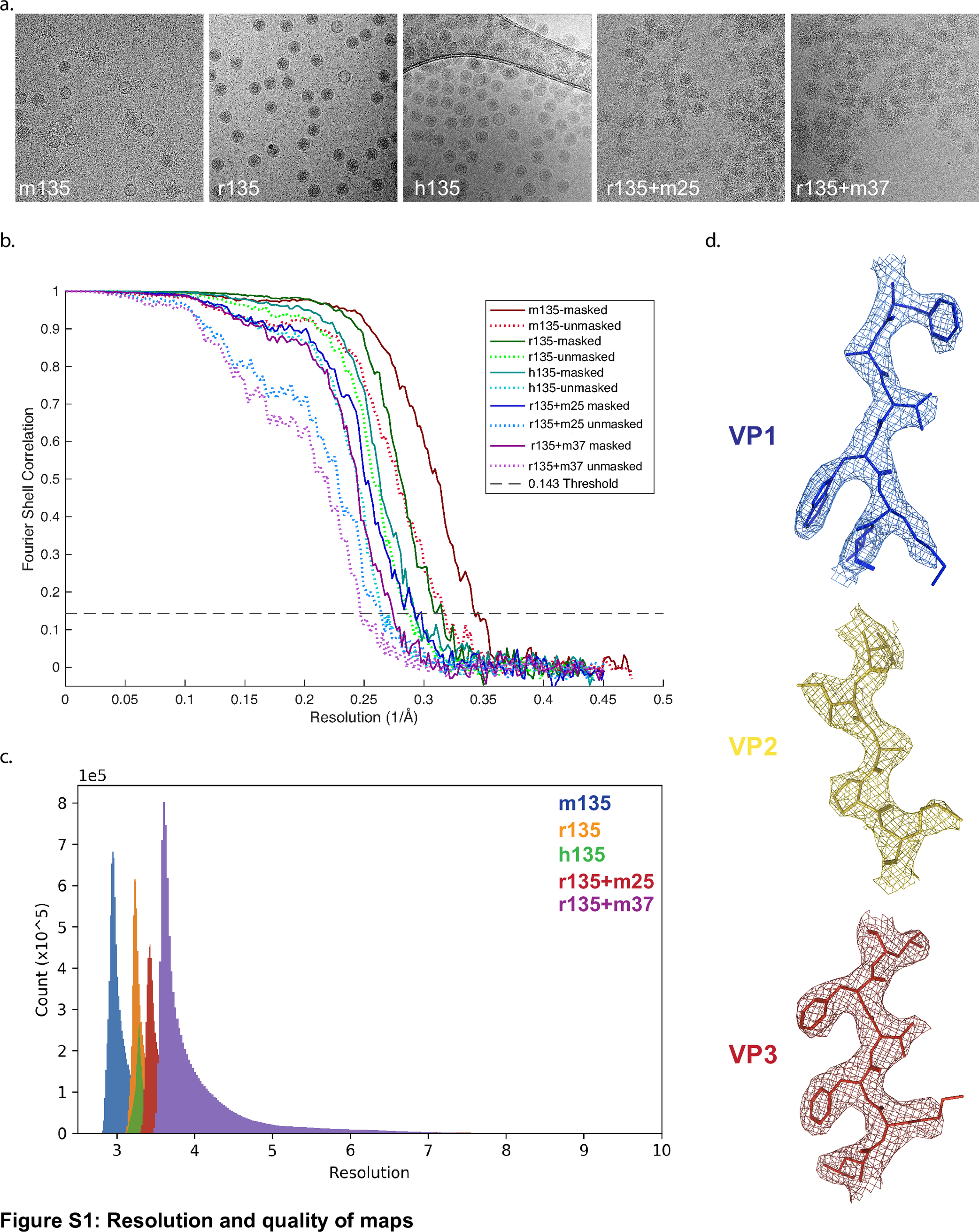

### Figure S2

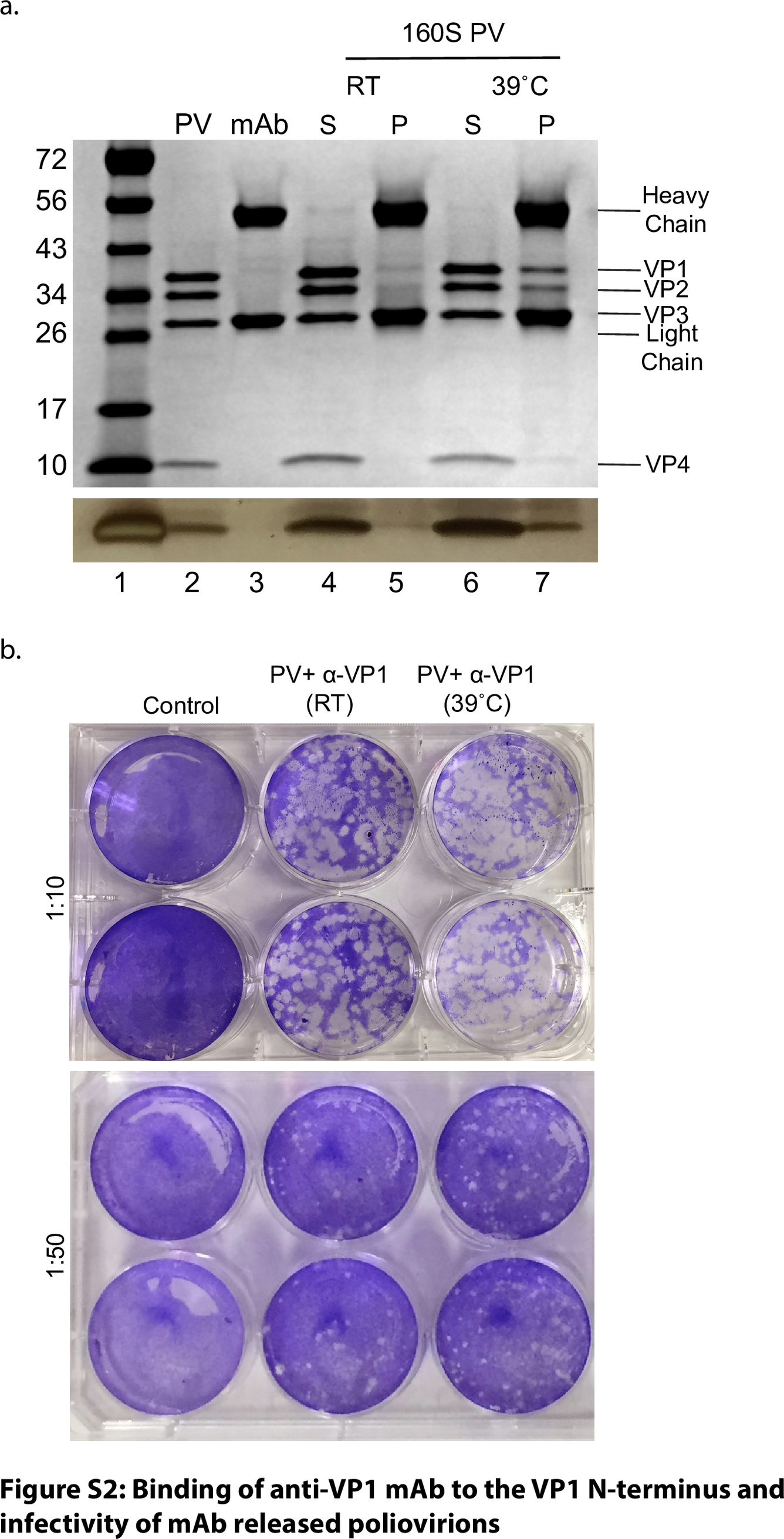

### Figure S3

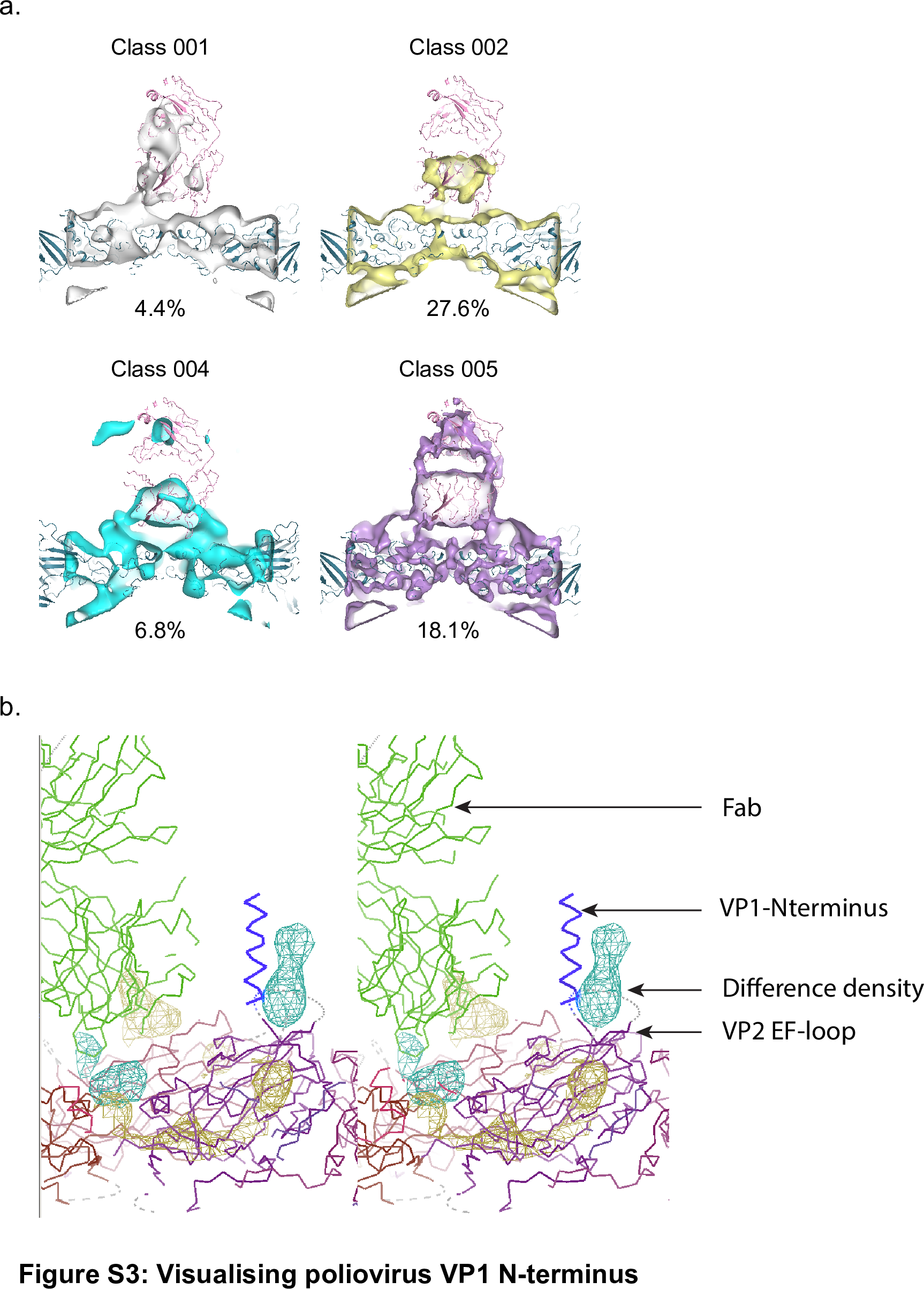

### Table S1

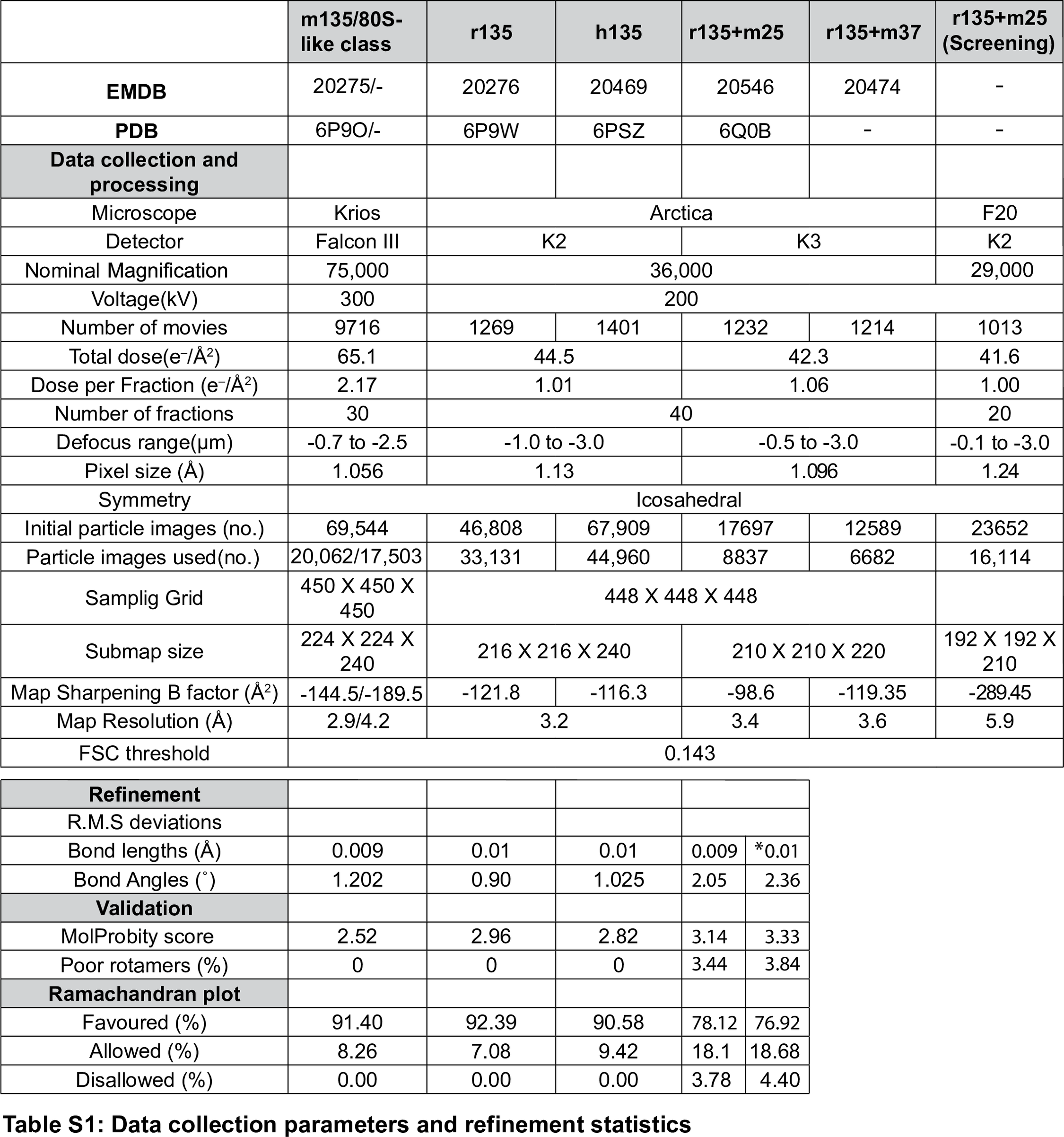

### Tablse S2

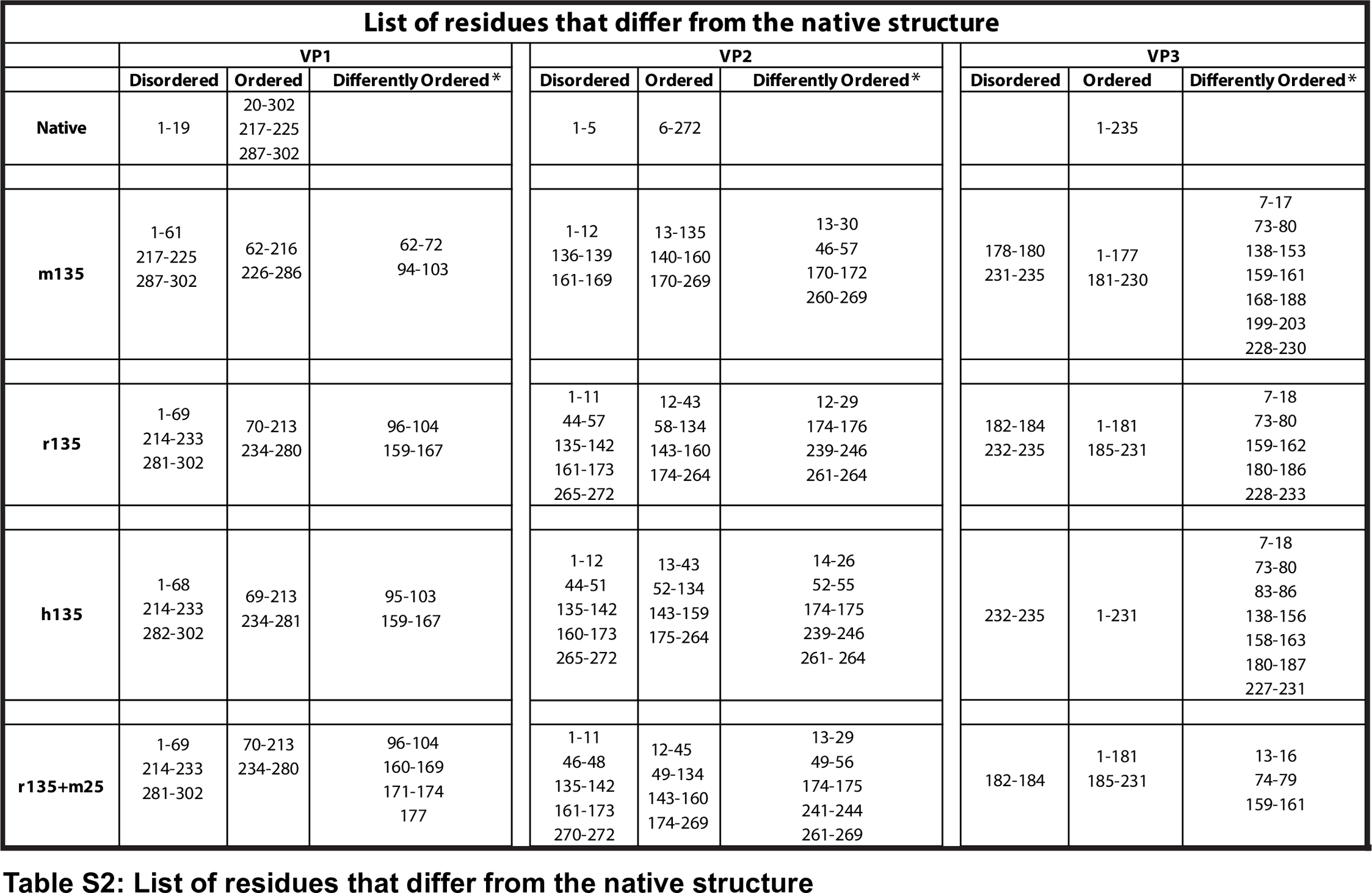
